# Lamin A/C Deficiency Enables Increased Myosin2 Bipolar Filament Ensembles Which Promote Divergent Actomyosin Network Anomalies Through Self Organization

**DOI:** 10.1101/2020.07.10.197731

**Authors:** O’Neil Wiggan, Jennifer G. DeLuca, Timothy J. Stasevich, James R. Bamburg

## Abstract

Nuclear envelope proteins influence cell cytoarchitecure by poorly understood mechanisms. Here we show that siRNA-mediated silencing of lamin A/C (LMNA) promotes contrasting stress fiber assembly and disassembly in individual cells and within cell populations. We show that LMNA deficient cells have elevated myosin-II bipolar filament accumulations, irregular formation of actin comet tails and podosome-like adhesions, increased steady state nuclear localization of the mechanosensitive transcription factors MKL1 and YAP, and induced expression of some MKL1/Serum Response Factor (SRF) regulated genes such as that encoding myosin-IIA (MYH9). Our studies utilizing live cell imaging and pharmacological inhibition of myosin-II, support a mechanism of deregulated myosin-II self-organizing activity at the nexus of divergent actin cytoskeletal aberrations resultant from LMNA loss. In light of our results, we propose a model of how the nucleus, via linkage to the cytoplasmic actomyosin network, may act to control myosin-II contractile behavior through both mechanical and transcriptional feedback mechanisms.

## Introduction

Mutations to LINC (linker of nucleoskeleton and cytoskeleton; Crisp et al., 2006) complex proteins or to proteins comprising the nuclear lamina, types A and B lamin intermediate filaments, give rise to pleiotropic human diseases collectively termed nuclear envelopathies (Burke and Stewart, 2006). The molecular and cellular pathogenesis of these diseases are not well understood but have been associated with perturbed gene regulation, altered cell mechanics and impaired force transduction between the nucleus and the cytoskeleton (Broers et al., 2004; Lammerding et al., 2004; Stewart-Hutchinson et al., 2008). Mechanical signals originating with cytoskeletal filaments are transmitted to the nucleus via the LINC complex (Wallrath et al., 2016). The mechanotransduction LINC apparatus consists of outer nuclear membrane nesprins, which bind cytoskeletal filaments and also couple to inner nuclear membrane localized Sun-domain proteins, which in turn are tethered to the nuclear lamina through association with lamins. Cells deficient of lamins A and C, products of a single gene via alternative splicing, or cells with LINC complex impairment, exhibit defects in cytoskeletal organization. However, there have been contrasting reports of cytoskeletal dysfunction following nuclear lamina disruption (via lamin A/C or LINC complex perturbations). Lamin A/C (LMNA) deficient cells, for instance, have variously been reported to have normal actin organization, increased stress fiber formation and contractile behavior, and alternatively, stress fiber loss with reduced contractility (Corne et al., 2017; Lammerding et al., 2004; Osmanagic-Myers et al., 2019; van Loosdregt et al., 2017). A rationale for this seemingly paradoxical collection of reports has not been identified.

Cytoarchitecture is directed by both F-actin arrangement and mechanical forces generated by the contractile activity of nonmuscle myosin-II (myo-II) molecular motors, which transport actin filaments relative to each other in an ATP-dependent cycle (Levayer and Lecuit, 2012). Individual myo-II motors are non-processive but form functional ensembles of variable numbers of bi-oriented units comprising single myo-II bipolar filaments. The activity of myo-II is regulated at multiple levels including phosphorylation by Rho-family effector kinases and through less characterized mechanisms such as via mechanical feedback from its own activity (Aguilar-Cuenca et al., 2014; Kasza and Zallen, 2011). Studies utilizing purified proteins in vitro, and of different cellular systems, have demonstrated the ability of myo-II to self-organize to diverse cytoskeletal patterns such as F-actin bundles, actomyosin asters and networks with a dynamic steady state (Backouche et al., 2006; Levayer and Lecuit, 2012). Significantly, myo-II activity can drive both the assembly and disassembly of networks of co-assembled F-actin and myosin.

Organization of the F-actin cytoskeleton significantly impacts nuclear function. For example, contingent upon F-actin assembly and mechanical signaling, key transcription co-activators such as YAP and MKL1 (also named MRTF-A and MAL) are either retained in the cytoplasm or shuttled to the nucleus, thereby modulating the expression of a broad array of genes important for varied cell functions (Aragona et al., 2013; Miralles et al., 2003; Olson and Nordheim, 2010). Recently it was suggested that F-actin cytoskeletal perturbations resulting from LMNA deficiency may be the result of abrogated actin-dependent signaling that prevents proper translocation of MKL1 to the nucleus (Ho et al., 2013b). Loss of MKL1 co-activation of a master cytoskeletal transcriptional module mediated by Serum Response Factor (SRF) and concomitant reduced expression of cytoskeletal genes, such as actin itself, was proposed as a mechanism for some cytoskeletal defects in LMNA deficient cells. Nevertheless, alternative mechanisms must still be explored since, for instance, nuclear translocation of MKL1 is not impaired in all LMNA deficient cells (Ho et al., 2013a). Furthermore, global reduced cytoskeletal gene expression appears insufficient to explain reported cases of enhanced contractile F-actin assembly in LMNA deficient cells.

To begin to elucidate the manner in which nuclear envelope proteins may affect cytoskeletal organization we have examined the effects of siRNA mediated depletion of LMNA in human cells. Remarkably, we find that depletion of LMNA, nesprins or Sun proteins similarly produce localized and contrasting actomyosin cytoskeletal perturbations, even within single cells. We identify that myosin-II assembly is enhanced in LMNA depleted cells. Our data link deregulated myosin-II and its self-organizing activity to both localized increased actomyosin assembly and disassembly associated with aberrant cytoskeletal phenotypes following LMNA depletion. We further demonstrate increased expression of myosin-2A in conjunction with elevated nuclear accumulation of both MKL1 and YAP, following LMNA depletion, in contrast to expectations based on some previous reports.

## Results

### Depletion of nuclear envelope proteins induces divergent actomyosin structural anomalies

Using multiple siRNAs, previously characterized to cause nuclear depletion of their respective targeted LINC complex proteins (Wiggan et al., 2017), we evaluated cytoskeletal organization following silencing of individual LINC complex proteins. Fascinatingly, silencing of LINC module components, (hereafter referring to either outer nuclear membrane nesprin1 or nesprin2, inner nuclear membrane Sun1 or Sun2 and LMNA), in multiple human cell types including HeLa, U2OS, non-transformed RPE-1, and normal fibroblasts (GM-2149) resulted in similar divergent aberrant actomyosin cytoskeletal phenotypes (Fig. 1). The majority of cells (∼55% for LMNA silencing) displayed increased numbers of thick basal stress fibers relative to controls. Additionally, we also observed two other general phenotypic groups. One group showed much reduced levels or complete absence of basal stress fibers (particularly fibers beneath the nucleus) and the other had cells with a disrupted network of stress fibers; and both were also characterized by the presence of actomyosin asters, F-actin condensates or fragmented F-actin bundles (for examples see enlargements Fig. 1A, D, H, G, N). Remodeling of the actomyosin network occurred throughout the entire cell and was not restricted to basal stress fibers. Notably however, actomyosin remodeling was regionally divergent such that individual cells simultaneously exhibited regions of increased F-actin bundling and those of stress fiber loss or fragmentation (e.g. Fig. 1J, arrows).

**Figure 1.**
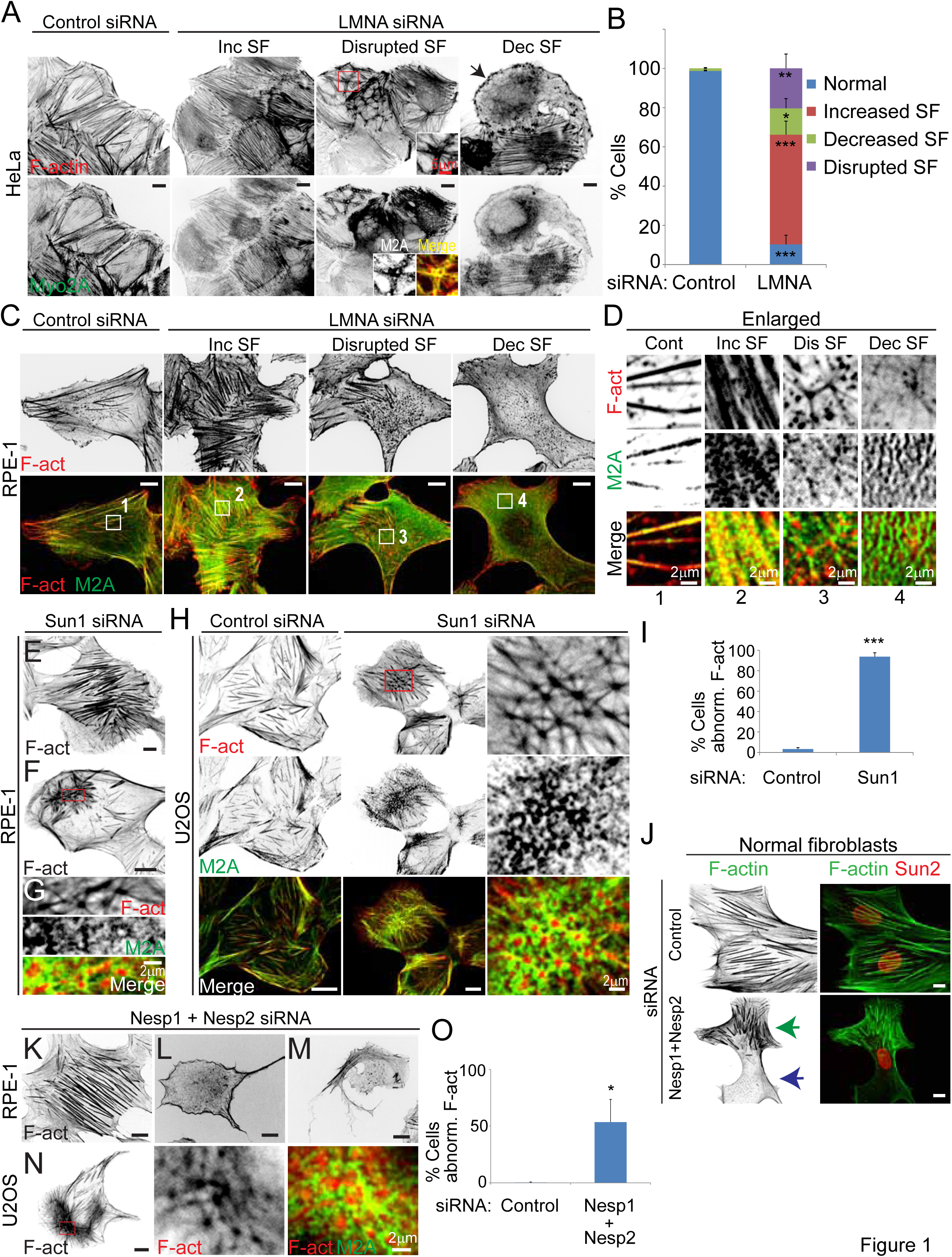
LINC complex disruption in multiple human cell types induces contrasting F-actin cytoskeletal anomalies. (A and B) Basal confocal immunofluorescence images of HeLa cells labelled for F-actin (phalloidin) and myosin-2A, and quantification of F-actin phenotypes. Arrow depicts cell with decreased numbers of stress fibers (SF) and the presence of F-actin condensates. Insets show enlargements and merged images (bottom row) of boxed region depicting rings of myosin-II surrounding interconnected F-actin asters. Values are mean ± SD, control n = 1192, LMNA siRNA n = 473. (C, D, E, F and G) Confocal immunofluorescence images of RPE-1cells siRNA treated for depletion of LMNA (C and D) and Sun1 (E-G). Numbered boxed regions (C, bottom row) are shown enlarged (D). Cells with increased SF (E), disrupted SF network (F), relative to control (C) are illustrated. Boxed region (F) is enlarged in G. (H) Immunofluorescence images of U2OS cells. Boxed region is enlarged in last column and shows a disrupted network of actomyosin asters. (I) Quantification of total abnormal F-actin phenotypes in RPE-1 cells, as classified in (B). Values are mean ± SD, control n = 180, Sun1 siRNA = 163. (J, K, L, M and N) Immunofluorescence images of normal human fibroblasts (J), RPE-1 (K-M) and U2OS (N). Increased SF formation (K), stress fiber loss (L and M), and disrupted stress fiber network (N) are shown. Green and blue arrows (J) depict regions of increased thick SF and stress fiber loss respectively. Boxed region (N) is enlarged in subsequent columns. (O) Quantification of total abnormal F-actin phenotypes in RPE-1 cells, as classified in (B). Values are mean ± SD, control n = 377, Nesprin1 (Nesp1)+Nesprin2 (Nesp2) siRNA = 256. Inc:increased, Dis:disrupted, Dec:decreased, F-act: F-actin, M2A: myosin-2A. * p < .05, ** p ≤ .01, *** p ≤ .001, ns, not significant, Welch’s t-test. Bars, 10 μm, except where noted.

Contrasting cytoskeletal phenotypes resulting from depletion of LINC module proteins could possibly result from different degrees of LINC protein silencing between cells. Opposingly, immunofluorescence staining of residual LMNA levels in both HeLa and RPE-1 revealed comparable low LMNA levels for cells displaying increased stress fibers to those exhibiting stress fiber loss (Fig. S1A, B). Further, quantification of basal F-actin accumulations revealed no significant correlations to residual cellular LMNA levels across cells. Curiously, heterogeneity in cytoskeletal defects within and between cells were highlighted features of fibroblasts from LMNA null mice where all cells had the same level of genetic LMNA depletion (Broers et al., 2004). Together, these data suggest that differing cytoskeletal phenotypes are not coupled to varying residual levels of LINC proteins following siRNA mediated depletion.

### LMNA silencing induces increased contractile cell behavior without significant changes to p-MLC levels

The predominance of cells with increased stress fiber formation following LMNA silencing suggested increased contractile myosin-II (myo-II) activity in response to this treatment. Myosin contractility is essential for focal adhesion maturation, which is associated with enhanced recruitment of core focal adhesion proteins (Schwarz and Gardel, 2012). Examination of focal adhesions revealed overall significant stronger accumulation (88% increase in RPE-1) of the focal adhesion protein paxillin to these sites in LMNA depleted cells for both HeLa and RPE-1 cells (Fig. 2A-C). Interestingly, as observed for stress fibers, individual LMNA depleted cells displayed heterogenous localized regions of either focal adhesion enhancement or reduction, coincident with that of stress fiber organization. Figure 2B, for example, illustrates a region of increased actomyosin and paxillin accumulations (region 1) in contrast a region of the same cell with stress fiber loss, which is associated with diminished paxillin accumulations (region 2). Heterogeneity in focal adhesion accumulations in LMNA silenced cells was also manifest by the broader distribution of adhesion intensities for these cells relative to controls (Fig. 2C). Given that myosin contractile activity can drive both actin network assembly and disassembly, the contrasting cytoskeletal rearrangements, evidenced by stress fibers and focal adhesions, are equally consistent with elevated contractile myo-II activity following LMNA disruption. We return to discuss the seeming paradox of these concurrent opposing responses in more details later.

**Figure 2.**
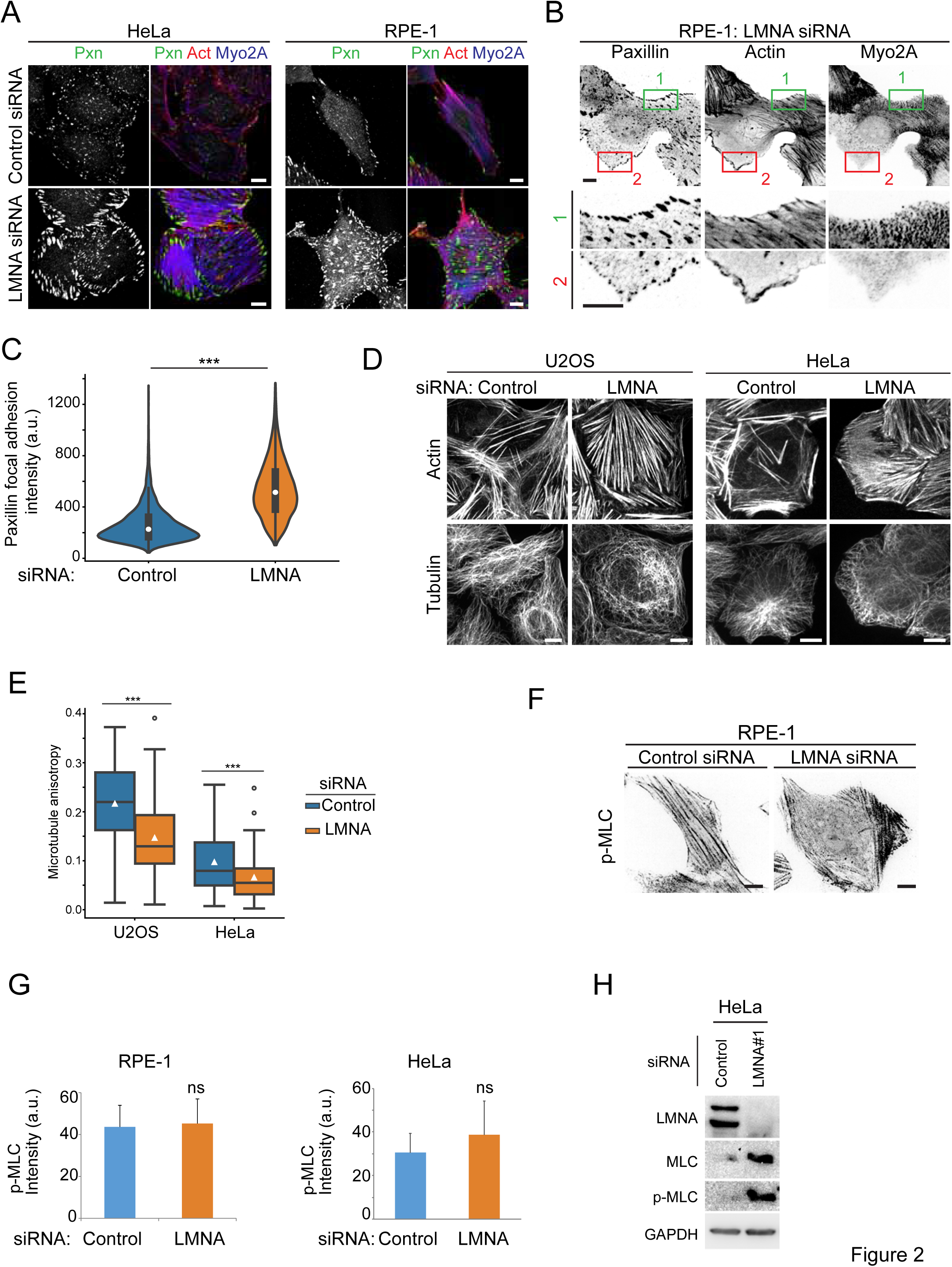
LMNA depletion elevates focal adhesion maturation without significantly altering global p-MLC levels. (A and B) Confocal immunoflouresence labeling for paxillin (pxn), F-actin (actin) and Myosin-2A (Myo2A). Boxed regions (B, upper panels) are labeled correspondingly in magnified lower panels. (C) Violin plot of focal adhesion paxillin intensity distributions for RPE-1 cells, n ≥ 30 cells/treatment. Lower and upper box horizontal lines of inner plot show 25^th^ and 75^th^ percentiles respectively, white circle shows the median value. (D and E) Immunofluorescence confocal images and quantification of microtubule anisotropy. Box plots (E) show interquartile range and white triangle is mean value. U2OS n ≥ 52, HeLa n ≥ 59. Immunofluorescence labeling (F), quantification of normalized cell p-MLC fluorescence intensity (G) and immunoblot (H) for active myosin-II (Ser19 phosphorylated myosin light chain). Values are mean± SD, RPE-1 n ≥ 101, HeLa n ≥ 426. Bars, 10 μm, except where noted. *** p < .001, ns, not significant, Welch’s t-test.

Microtubule disassembly is known to stimulate formation of myo-II mini-filaments, stress fibers and focal adhesion (Krendel et al., 2002; Liu et al., 1998; Rafiq et al., 2019) so we assessed microtubule organization in different LMNA depleted cell types. HeLa and U2OS LMNA silenced cells displaying increased stressed fibers both maintained microtubules similar to controls (Fig. 2D). Nevertheless, LMNA depletion induced microtubule reorganization, from a more ordered radial pattern in control cells, to overlapping curved bundles manifested by increased microtubule fibrillar disorder (Fig. 2E, reduced anisotropy, where 0 is isotropic and 1 is perfectly ordered; Boudaoud et al., 2014). These data show that LMNA deficiency contributes to altered microtubule patterning, however, the absence of gross microtubule loss in LMNA depleted cells suggests that the latter feature was not a significant mechanistic factor to increased actomyosin assembly.

We next examined whether increased contractile behavior following LMNA depletion was associated with altered myo-II phospho-activation. Immunostaining for S19 phosphorylated myosin light chain (p-MLC, phospho-active myosin), revealed a trend towards increased p-MLC, however, p-MLC levels were not significantly different between controls and either RPE-1 or HeLa LMNA depleted cells (Fig. 2F-G). Likewise, whereas immunoblots indicated increased (∼ 4-fold) p-MLC levels for LMNA depleted cells, this rise was in parallel to a ∼5-fold increase in total MLC levels, relative to controls (Fig. 2H). Collectively, these data demonstrate that LMNA depletion stimulates increased cell contractile activity without apparent significant increases to steady state p-MLC levels.

### LMNA deficiency induces aberrant membrane associated actomyosin remodeling

Myosin activity is important for dynamic remodeling of actin filaments in organization of various membrane structures (Koster et al., 2016; Vogel et al., 2017). If LMNA contributes to altered myo-II function, we hypothesized that LMNA deficiency may induce perturbations to membrane associated actomyosin. In vitro, myosin self-organization can produce aster-like structures, where myo-II localizes to center of the aster (Backouche et al., 2006; Soares e Silva et al., 2011), in contrast, aster-like structures at the basal plasma membrane in LMNA depleted cells displayed a central core of actin surrounded by a ring of myo2A (see for example Fig.1A, inset; 1D, Dis SF). The latter organization is reminiscent of podosomes, plasma membrane-extracellular matrix adhesion structures, found in certain normal cell types and some transformed cells (Alonso et al., 2019; Marchisio, 2012). The actin core of podosomes is surrounded by outer rings of integrin-associated proteins, such as paxillin, and myo2A respectively. Confocal images resolved below the diffraction limit using super-resolution radial fluctuations (SRRF) computation (Gustafsson et al., 2016) revealed this organization among actin condensates in LMNA depleted HeLa cells (Fig. 3A-B). Unlike HeLa cells, podosome-like structures in RPE-1 cells did not assemble a ring of paxillin (Fig. S2), suggesting a difference between these two cell types in their ability to form full podosome-type adhesions. Other distinctive features of podosome-type adhesions, such as recruitment of the actin binding protein cofilin to the actin core (Yamaguchi et al., 2005) was evident for podosome-like adhesions in HeLa cells (Fig. 3C). Myosin contractile activity is essential for oscillatory intensity fluctuations of the actin core of podosomes (van den Dries et al., 2013) and this oscillatory behavior was displayed by podosome-like adhesions that formed in LMNA deficient HeLa cells (Fig. 3D, Video S1). Live imaging suggested spontaneous formation of the actin core, which appeared to intensify and stabilized by mutual assembly reinforcement with proximal encircling myo2A puncta (Fig. 3E). These data highlight cell-type specific aberrant membrane associated actomyosin remodeling in association with LMNA deficiency.

**Figure 3.**
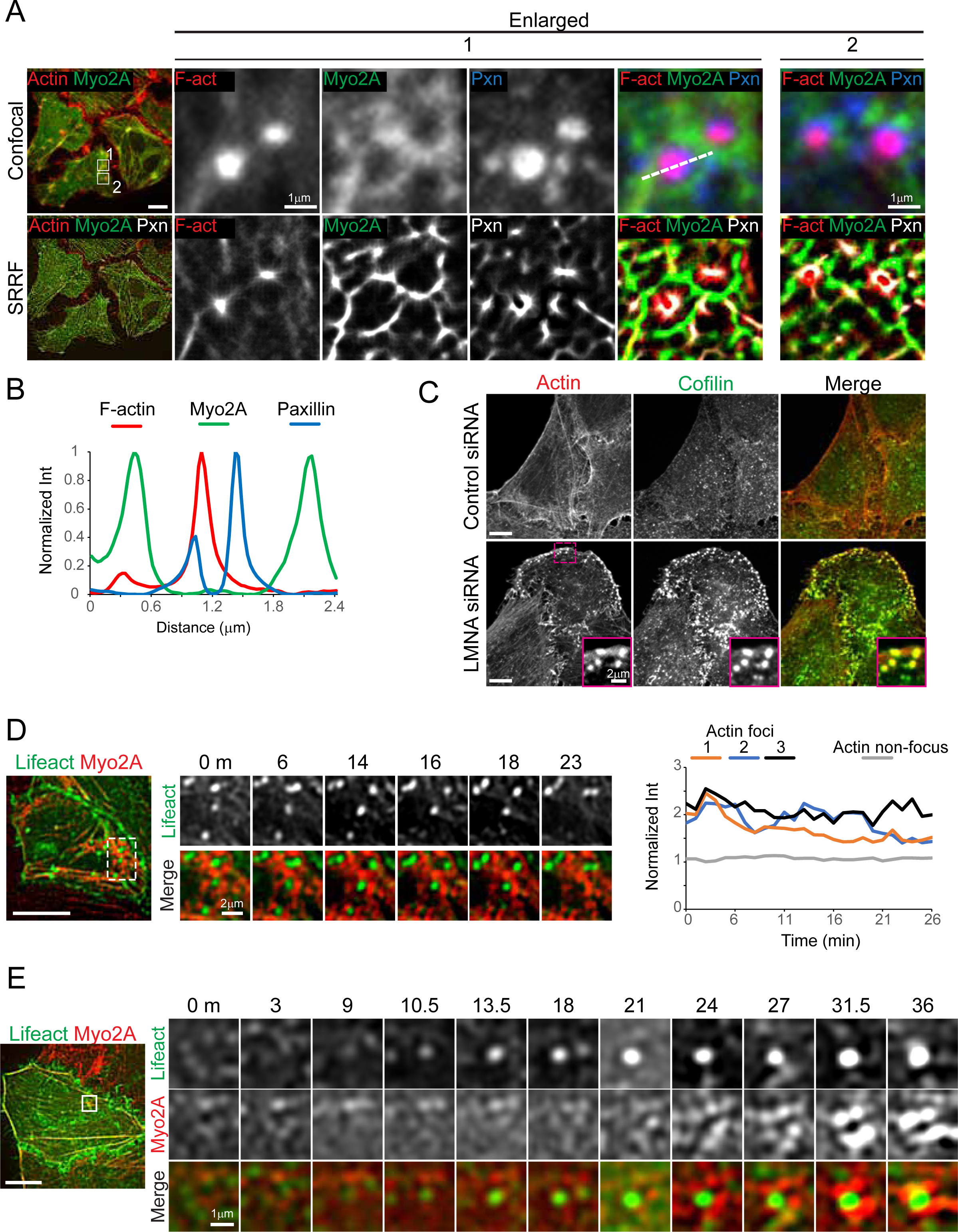
LMNA silencing induces formation of podosome-type adhesions. (A) Confocal immunofluorescence (upper panels) and SRRF super-resolved images of LMNA siRNA treated HeLa cells labeled for paxillin (pxn), F-actin (F-act) and myosin-2A (myo2A). Numbered boxed regions (left column) are shown enlarged, correspondingly. (B) Normalized line intensity profile plots, from line shown for region 1, top row, obtained from SRRF images. (C) Immunolabeled confocal images of fixed HeLa cells. Boxed region (LMNA siRNA) is shown magnified in insets. (D and E) Confocal video stills of HeLa Myo2-mCh cells, coexpressing GFP-Lifeact for F-actin labeling and siRNA treated for LMNA silencing. Boxed regions (left panels) are enlarged in time-series images (right panels). See also Video S1. Graph (D) plots podosome-like structure actin core intensity with time. Scale bars, 10 μm.

Live cell analyses of both HeLa and RPE-1 cells showed an increased presence of dynamic membrane linked actin structures known as comet tails (Cameron et al., 2000), in LMNA depleted cells relative to controls (Fig. 4A, Videos S2-3). In vitro models of actin comet formation do not show necessity for myosin, however, some reports implicate association of myo-II with comet activity in cellular models of pathogen transmission (Lum and Morona, 2014; Rathman et al., 2000). We assessed whether myo-2A was associated with these actin assemblies in LMNA silenced cells. As illustrated, Fig. 4B-D (arrowheads), we identified several instances where translocation of actin-myo2A structures to a cluster was concurrent with the formation of an actin focus or comet. Figure 4D (yellow arrow, and see Video S3) also shows that myo2A can incorporate to comet tails, further suggesting myo-II involvement in the formation of these structures in LMNA depleted cells. Altogether, our analyses thus far are consistent with a hypothesis that deregulated actomyosin remodeling contributes to the major actin cytoskeletal defects, including those associated with cell membranes, obtained from LMNA deficiency.

**Figure 4.**
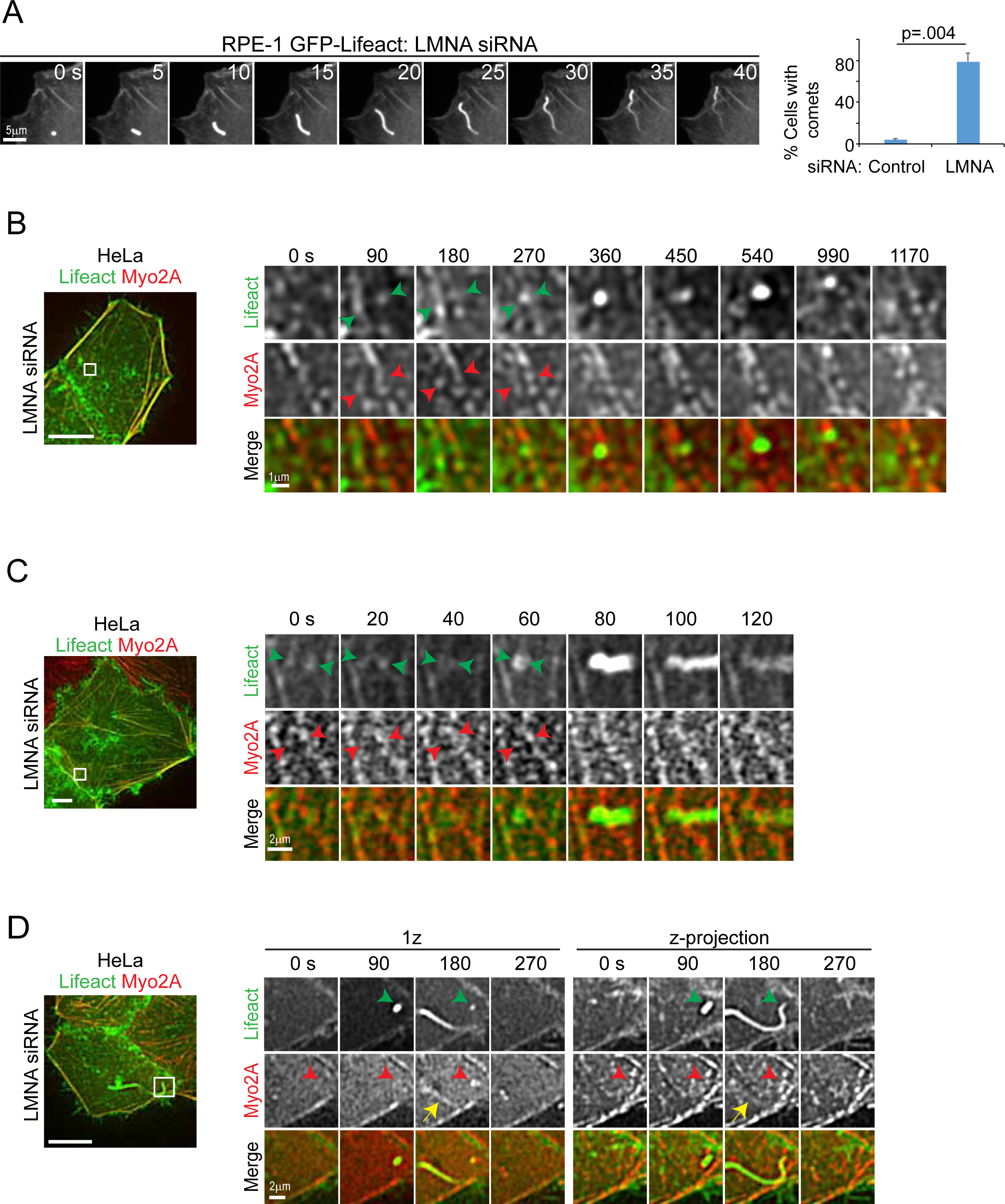
Localized actomyosin remodeling is associated with increased assembly of F-actin to comet tails in LMNA depleted cells. (A) Confocal time-lapse series and quantification of F-actin comet formation in RPE-1 cells. Values are mean ± SD, n = 90/treatment. Welch’s t-test. See also Video S2. (B, C and D) Confocal video stills of HeLa Myo2-mCh cells, coexpressing GFP-Lifeact for F-actin labeling and siRNA treated for LMNA silencing. Left panels are basal cell z-projections and boxed regions are enlarged in time-series of selected z-planes (right panels). Arrowheads (B, C) depict movements that cluster actin-myosin structures at sites of heightened actin assembly. Arrowheads and yellow arrow (D) show myo2A accumulation (red arrowhead) at a site of comet tail formation (green arrowhead) and myo2A localization to the nascent comet tail (yellow arrow), respectively, for single and z-projected images. See also Video S3. Scale bars, 10 μm, except where noted.

### Increased nuclear accumulation for mechanosensitive transcription factors MKL1 and YAP in LMNA depleted cells

Our results indicate that LMNA deficiency caused increased myo-II contractile behavior resulting, for example, in increased stress fibers and focal adhesion maturation. These data insinuated that LMNA deficiency may in fact stimulate mechanosensitive transcription pathways, such as for MKL1 and YAP. Furthermore, recent studies implicate mutual dependence of both MKL1 and YAP for regulation of their respective transcription targets (Foster et al., 2017). Examination of MKL1 and YAP cellular localization showed their increased nuclear localization following LMNA silencing for both HeLa (Fig. 5A) and RPE-1 cells (Fig. 5B-D) grown in normal serum-containing medium. Our results for MKL1 in cells grown in normal growth medium differ from those previously reported for some LMNA deficient cells stimulated from quiescence, where nuclear MKL1 accumulation was abrogated relative to controls (Ho et al., 2013b). However, at conditions of previous reports, i.e. serum stimulation from quiescence, we did not detect any significant differences in MKL1 nuclear accumulation in LMNA depleted cells relative to controls (Fig. 5B-C). These data imply that effects of LMNA deficiency on MKL1 nuclear translocation maybe contextual. Separately, we uncovered strong correlations (Pearson’s r ≥ 0.8) between the nuclear/cytoplasmic ratio for MKL1 relative to those of both G-actin (Fig. 5F) and YAP (Fig. 5G), consistent with likely coordinate regulation of MKL1 localization with both factors. Interestingly however, MKL1 nuclear/cytoplasmic distribution did not show a strong correlation to cellular G/F-actin ratios (Fig. 5E), as might be expected based on current models of MKL1 activation. Altogether, these results advocate that LMNA depletion induced activation of both MKL1 and YAP nuclear translocation in the context of steady state cell growth conditions.

**Figure 5.**
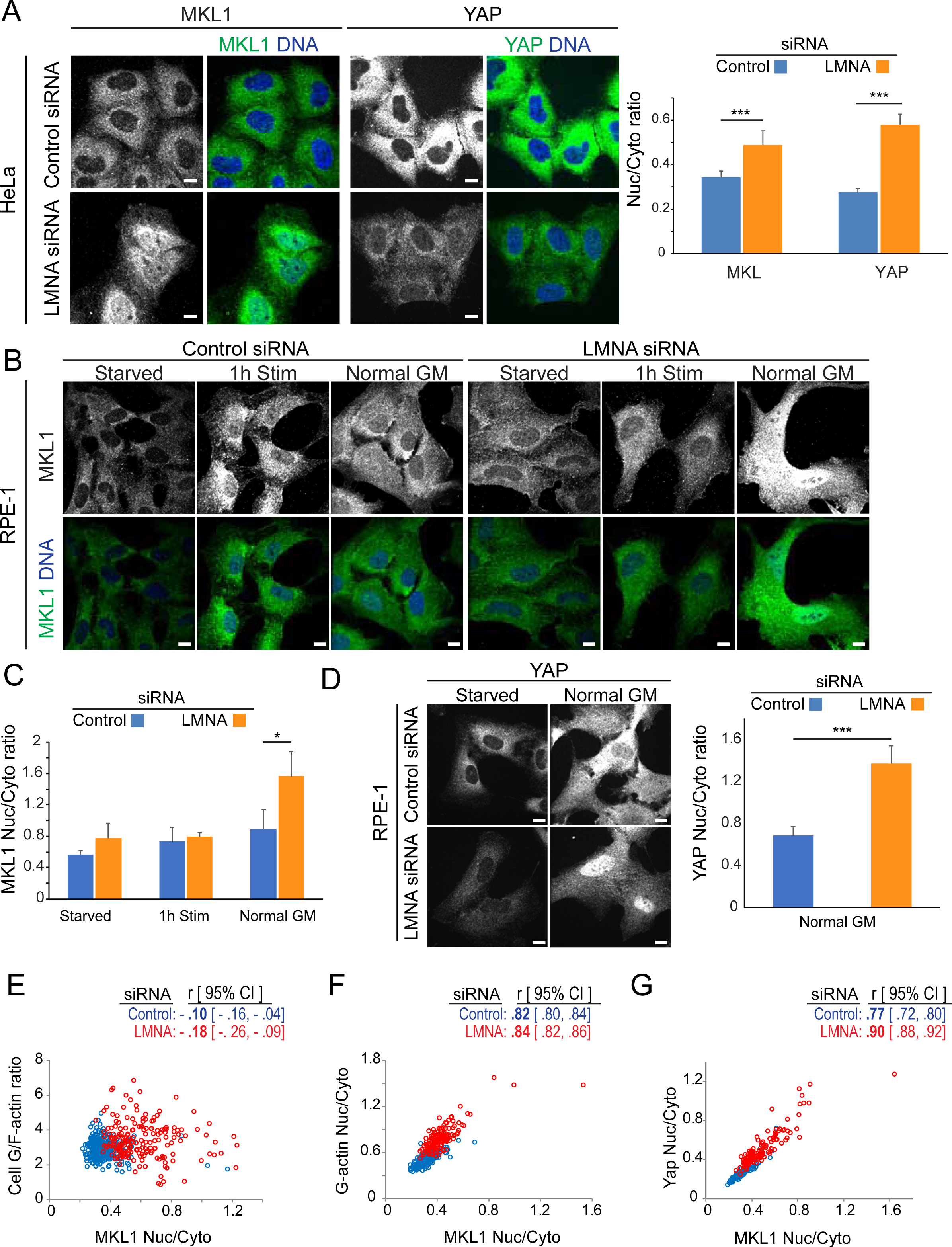
LMNA silencing induces nuclear accumulation of mechanosensitive transcription factors. (A, B, C and D) Representative nuclear level confocal immunofluorescence images and quantification of transcription factor nuclear-cytoplasmic ratios for HeLa cells (A) or RPE-1 cells (B, C, D) at 72h post siRNA treatments, as indicated. Cells (A, and as specified) were cultured in normal growth medium (Normal GM) or serum stimulated (1h stim) from serum starvation (starved) as indicated. Graph values are mean ± SD, n ≥ 115/treatment. (E, F and G) Representative per cell normalized fluorescence intensity correlation plots of HeLa cells treated as indicated. Values, r, are mean Pearson correlation coefficient and [95% confidence interval], n ≥ 283/treatment. Nuc:nuclear, Cyto:cytoplasmic. * p ≤ .05, *** p ≤ .001, Welch’s t-test.

In light of observed increased nuclear MKL1 accumulation in LMNA deficient cells, we examined whether known MKL1/SRF target genes were responsive to MKL1 regulation following LMNA depletion. Ectopic expression of two different constitutively active MKL1 mutant proteins (Flag-MKL1-N100, lacks an N-terminal actin binding motif required for its G-actin dependent cytoplasmic retention; and Flag-MKL1-STS/A, S449, S454, and T450 mutated to A, is refractory to ERK regulated nuclear export; Muehlich et al., 2008), showed both nuclear and cytoplasmic localization (Fig. S3), as observed by others (Willer and Carroll, 2017).

Significantly, active MKL1 mutants promoted expression of SRF target genes, for both control and LMNA silenced cells, without mitigation of cytoskeletal anomalies resulting from LMNA silencing (Fig. 6A, S3). For SRF targets assessed, LMNA silencing led to increased transcript levels for ACTB and MYH9, however, VCL levels were reduced by five-fold (Fig. 6A). These results suggest differential and complex regulation of MKL1/SRF target genes in response to LMNA deficiency.

**Figure 6.**
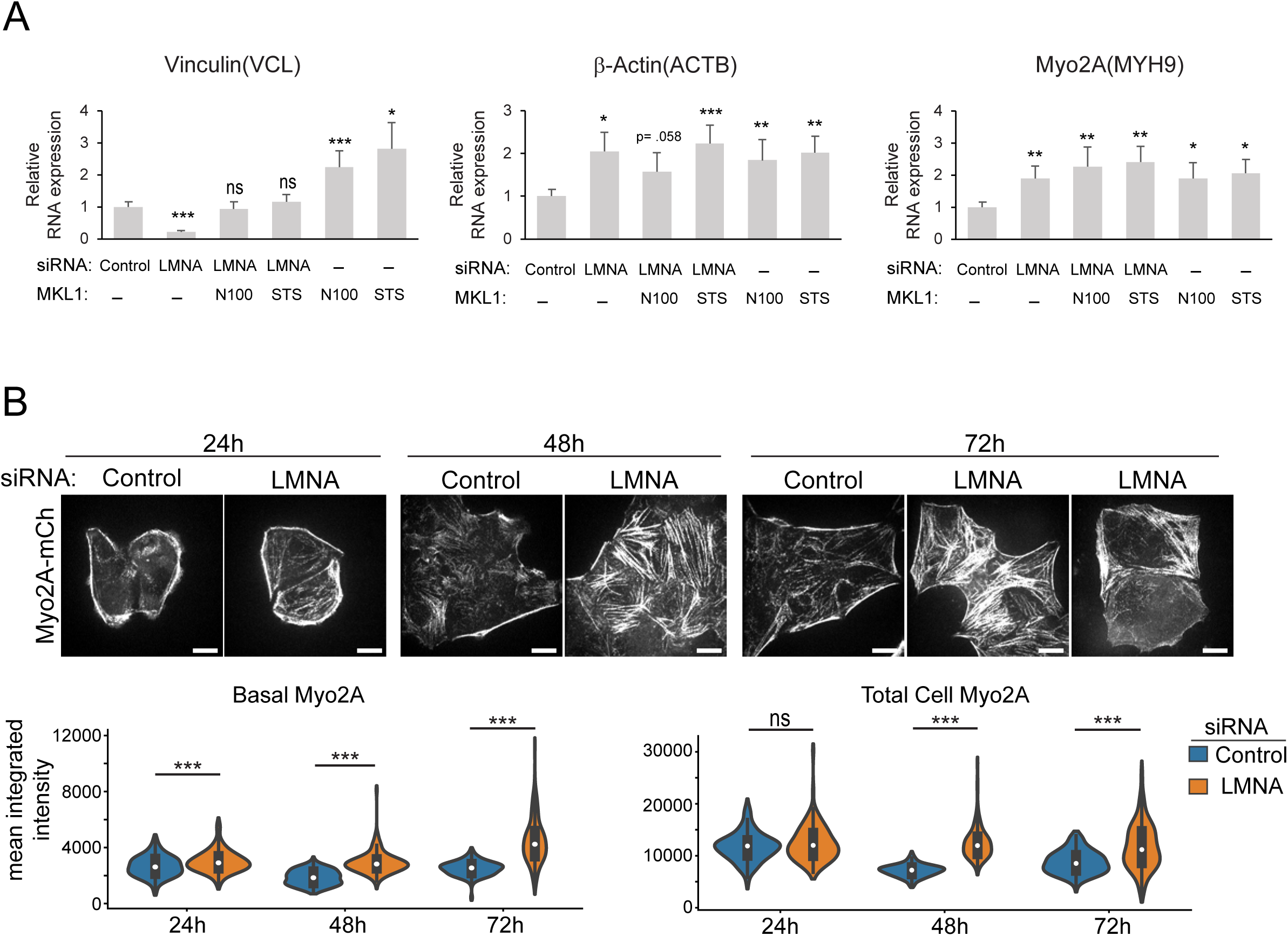
LMNA silencing alters MKL1/SRF target gene expression. (A) Quantitative PCR analysis of MKL1/SRF target gene expression, normalized to GAPDH, in HeLa cells treated as indicated and assessed at 72 h post siRNA treatment. Values are mean ± SEM, n=4. (B) Live cell fluorescence confocal images and fluorescence quantification of endogenous mCherry tagged myo2A (Myo2A-mCh) in HeLa cells at times post siRNA treatment, as indicated. Basal myo2A corresponds to measurements from basal confocal planes and total cell to the entire z-volume. *** p ≤ .001, ns, not significant, Welch’s t-test; n ≥ 96/treatment. Bars, 10 μm.

Congruent with increased myo2A transcript levels as measured at 72h post siRNA treatment (Fig. 6A), myo2A protein levels increased progressively following LMNA silencing (Fig. 6B). Myosin-2A fluorescence intensity measurements from cells expressing mCherry-tagged myo2A from the endogenous locus (HeLa Myo2A-mCh) provided a direct readout of myo2A protein levels throughout the period of LMNA silencing in single population of live cells. By this procedure increased myo2A protein accumulation, both basally and cell-wide, was evident for LMNA depleted cells by 24h post siRNA treatment and this was maintained to 72h post treatment, relative to controls (Fig. 6B). Our live cell studies also suggested progressive development of cytoskeletal anomalies in LMNA depleted cells with the presence of increased stress fibers becoming pronounced by 48h post siRNA treatment, whereas cells with central stress fiber loss or a disrupted network were more evident at 72h (Fig. 6B). Altogether, these data support activation of a contractile regulatory cascade entailing stimulation of mechanosensitive transcription factors with feedback to augmented myo2A expression.

### Increased Myosin-II bipolar filament assembly in LMNA depleted cells

Since our earlier results suggested that LMNA depletion may be associated with elevated myo-II contractile activity in the absence of significant increases to p-MLC levels, we interrogated other possible mechanisms linked to increased myo-II contractility. Computational models predict sharp sensitivities of myo-II force generation to bipolar filament ensemble size (Stam et al., 2015). Myosin-2A formed punctate structures in both fluorescent images of fixed cells labeled by antibodies to the C-terminal tail (e.g. Fig. 1D), or live cells expressing C-terminal mCh-tagged myo2A. From SRRF super resolved live cell images where endogenous myo2A was labeled by both N-terminal GFP and C-terminal mCh, via CRISPR/Cas9 gene editing (Fig. 7A), we determined that individual C-terminal myo2A labeled punctum match the profile of single bipolar filaments. As expected, C-terminal punctum were flanked on either side by spots corresponding to N-terminal head motifs. Accordingly, gaussian fits to intensity line scans of dual labeled myo2A puncta showed a mean bipolar filament length of 334 nm (n=7, Fig. 7A), in line with previous measurements made from EM ultrastructure studies (Niederman and Pollard, 1975). Measurements along stress fibers demonstrated more intense accumulation of myo2A to puncta for both HeLa and RPE-1 cells (Fig. 7C, D, F, G). Myo2A-mCh peak intensities, a reflection of the quantity of myo2A motors in each bipolar filament or punctum, were on average 53% greater for LMNA depleted HeLa cells (Fig. 7B), indicating larger bipolar filament ensembles for this condition relative to control.

**Figure 7.**
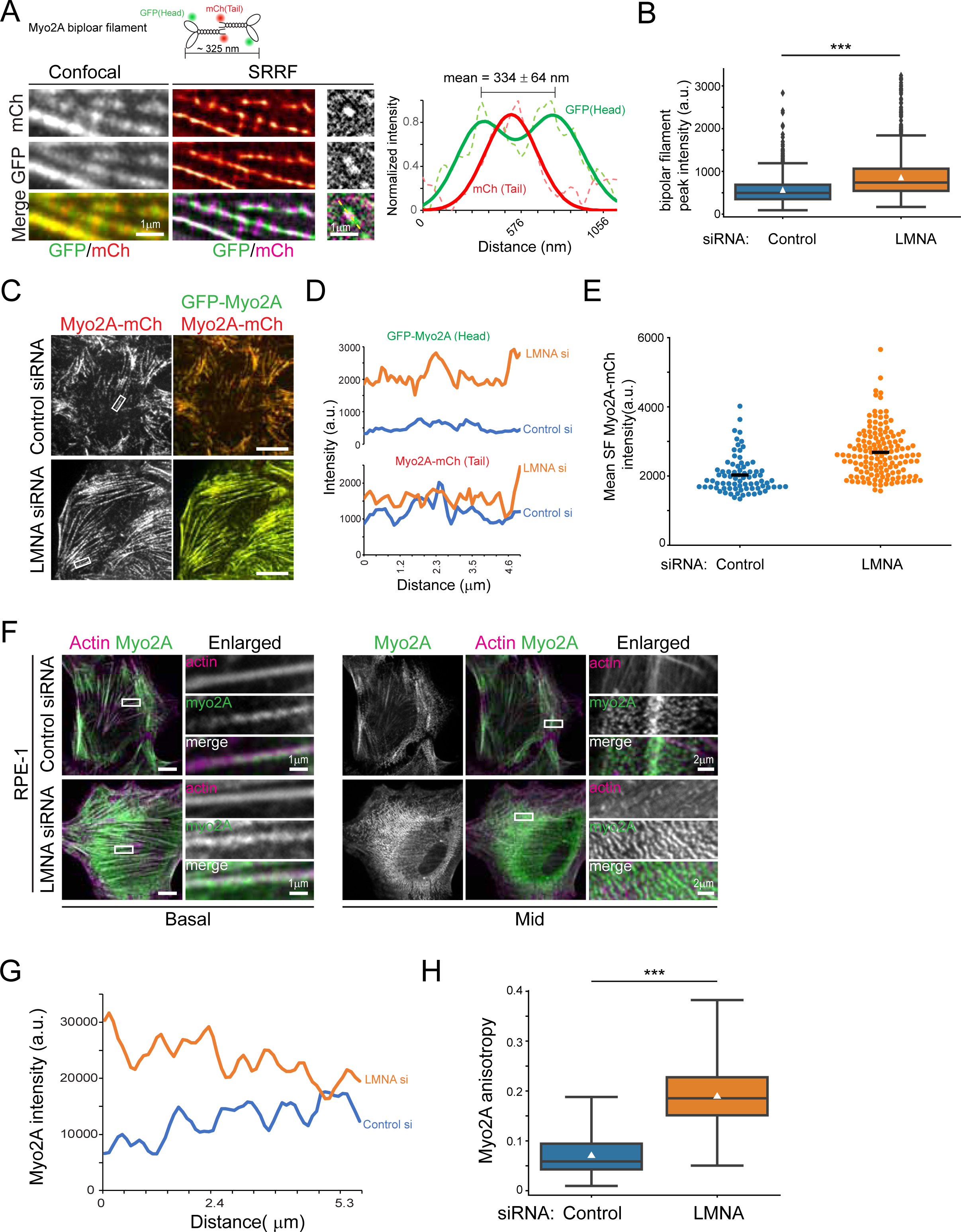
Increased myosin-II bipolar filament assembly in LMNA depleted cells. (A) Confocal and SRRF images of myosin-II bipolar filament organization in HeLa cells expressing dual tagged (as illustrated in cartoon) endogenous myo2A. Graph shows line intensity profile (marked by white line in right column) of an isolated bipolar filament. Plots (dashed lines) and gaussian fits (solid lines) are shown. (B) Plots of bipolar filament peak intensities as obtained from mCh (myo2A tail) images, n ≥ 968/treatment. (C and D) Basal confocal fluorescence images and line intensity profile plots of stress fibers shown in boxed region (C) for dual labeled myo2A in HeLa cells. (E) Quantification of myo2A intensity along stress fibers assembled following blebbistatin treatment and washout, n ≥ 78/treatment. Horizontal lines are mean values. (F) Basal and mid-cell level confocal immunofluorescence images of RPE-1 cells. Boxed regions are shown in enlargements. (G) Myo2A line intensity plots along stress fibers shown in basal boxed regions (F). (H) Quantification of myo2A filament anisotropy, n ≥ 83/treatment. White triangles are mean values. Scale bars, 10 μm or as indicated. *** p ≤ .001, Welch’s t-test.

In addition to the increase of myo2A to basal stress fibers, for RPE-1 LMNA depleted cells there was also pronounced accumulation and ordering of myo2A to elongated arrays of bipolar filaments in features previously characterized as myosin ‘ribbons’ or ‘stacks’ (Fig. 7F, (Fenix et al., 2016; Verkhovsky et al., 1995). Measurements of the fibrillar ordering of myo2A, affirmed augmented anisotropy for LMNA silenced RPE-1 cells relative to controls (Fig. 7H). In contrast to myo2A bipolar filaments in stress fibers, myo2A stacks concentrated at mid to apical regions of cells and did not appear to be associated with contractile actin structures, instead, these stacks were arranged as orthogonal arrays to sheets of thin of F-actin bundles (Fig. 7F). Our data here establish that LMNA deficiency results in enhanced myo-II bipolar filament assembly and suggest the involvement of cell type specific variables in accompanying filament stack expansion.

### Stress fiber disorder in LMNA deficient cells requires myosin-II contractile activity

To further define the requirement for myosin-II in the induction of contractile structures such as stress fibers in LMNA depleted cells, we evaluated whether myo-II inhibition altered assembly of such features. HeLa Myo2A-mCh cells were siRNA co-treated with blebbistatin (Straight et al., 2003), an inhibitor of myo-II which blocks motor ATPase activity and maintains myosin in a weak actin binding state, followed by live cell image analysis. Blebbistatin (blebb) treatment resulted in robust stress fiber disassembly for both LMNA silenced and control cells (Fig. 8A, S4A, Video S4). Strikingly however, whereas only 6% of control cells had any remaining myo2A labeled stress fibers at 2h post blebb treatment, 47% of LMNA silenced cells displayed the persistence of some stress fiber labeling at this time point (Fig. 8A, B, S4A, red arrows). Thus, while myo-II activity is required for the presence of increased stress fibers in LMNA depleted cells, these data also provide that LMNA deficiency results in stronger myo-II-actin interaction that is more resistant to blebb inhibition. Consistently, within 2h following washout of blebb, a significantly greater percentage of LMNA silenced cells had developed increased numbers of thick stress fibers relative to control cells (Fig. 8A, C, S4A) and these stress fibers accumulated more myo2A (32% increase, p < .0001, Welch’s t-Test; Fig. 7E), in comparison to those formed in control cells. Of note, following blebb washout, as cells re-assembled stress fibers, LMNA depleted cells were more prone to contract and undergo apoptosis relative to control cells (Fig. 8D, S4B). This latter result suggested greater sensitivity to mechanical stress for these LMNA deficient cells.

**Figure 8.**
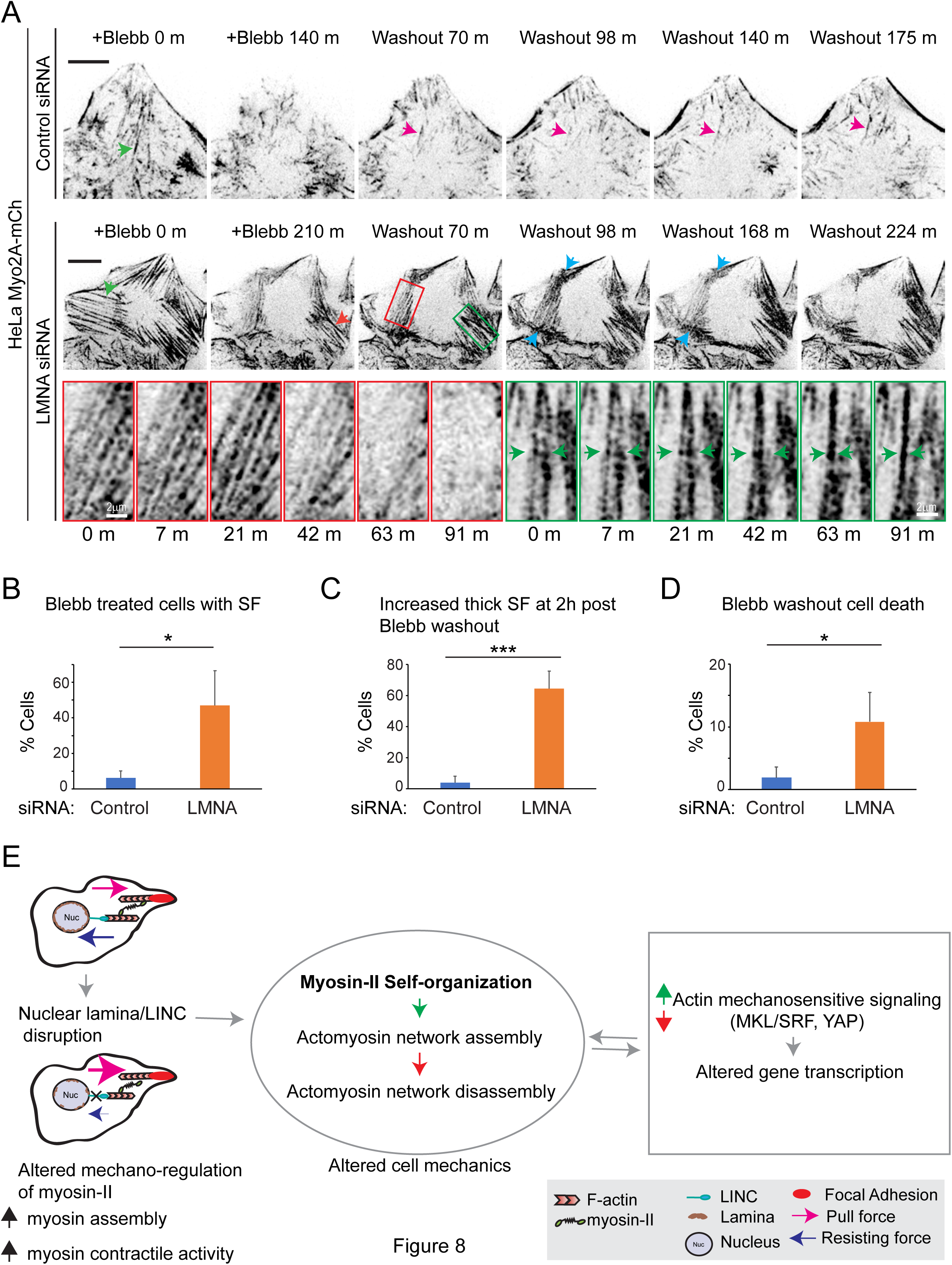
Myosin-II contractile activity instigates localized dynamic actomyosin remodeling in LMNA deficient cells. (A) Fluorescence time-lapse series of blebbistatin treatment (+Blebb) and drug washout. Green arrowheads depict stress fibers (SF) which disassembly during bleb treatment. Red arrowhead shows SF which persists at 2h post blebb treatment. Magenta arrowhead illustrates dynamics of a new, post-drug washout SF for a control cell. Red and green boxed regions (LMNA siRNA, upper panels) are magnified in lower panels and highlight localized actomyosin remodeling to SF disassembly and SF bundling, respectively, at ∼ 1h post blebb washout. Blue arrowheads depict regions of apparent focal myosin contractile activity. Green arrowheads depict merger of two adjacent SFs. See also Video S4. (B, C and D) Quantification of cells with SFs at 2h post blebb treatment (B), Quantification of increased thick SF assembly at 2h post blebb washout (C) and Quantification of cell death occurring within 3h post blebb washout (D). Values are mean ± SD, n ≥ 242/treatment. * p ≤ .05, *** p ≤ .001; Welch’s t-test. (E) Proposed mechanistic model for actomyosin organization dysfunction in response to disruption of the nuclear LINC module.

Finally, we assessed signs of myo-II self-organization to different patterns from live cell time series images, following blebb washout. Myo2A structures were dynamic for both control and LMNA depleted cells following blebb removal. For example, Figure 8A (control siRNA, magenta arrow) illustrates fluctuations in the organization of a newly formed stress fiber in a control cell, and that this stress fiber persisted in some form for over an 1h. For LMNA depleted cells we also detected dynamic myo2A behaviors which apparently contributed to simultaneous localized stress fiber bundling (Fig. 8A, green boxed region) and stress fiber disassembly (red boxed region) within a single cell. Stress fiber bundling looked to entail the translocation of adjacent stress fibers and their merger into a single thicker bundle (Fig. 8A, green arrows), concomitant with the merger of myo2A puncta to more intense filaments. Localized stress fiber disassembly in this instance seemed to involve opposing contractile forces focused at either ends of the stress fiber bundles (Fig. 8A, blue arrows). Both forms of myo-II patterning behaviors occurred over a period of tens of minutes, suggesting a relatively non-rapid time scale for myo-II self-organization in this cellular context. Combined, our live cell analyses support a pivotal role for deregulated myo-II motor activity and consequent exacerbated myo-II self-organizing behaviors in the etiology of actin cytoskeletal disorganization resulting from LMNA deficiency.

## Discussion

The opposing actin cytoskeletal alterations we observe in single populations of cells, following LINC module perturbations, have been reported independently as distinct aberrations by previous seemingly conflicting reports. In our experimental system a simple explanation may have been that different residual levels of LMNA depletion induced different cytoskeletal anomalies. However, the totality of our data, including Fig. S1, is unsupportive of this explanation. Furthermore, this principle cannot account for similar heterogenous cytoskeletal abnormalities reported within individual and across a population of germline LMNA knock-out cells (Broers et al., 2004). Blending our data with those from other studies we offer a rationale for the superficial paradox. We conclude that divergent actomyosin phenotypes are the result of a common process, myosin-II self-organization. Differing actomyosin arrangements within and across cells represent varying phases of a dynamic self-organizing contractile process that can transition between localized states of actomyosin assembly and disassembly. Our showing of increased presence of stress fibers, focal adhesions and actomyosin aster-like structures following LMNA depletion in various cell types, are all consistent with elevated myo-II contractile activity. Less appreciated, but anticipated based on numerous prior studies, actomyosin disassembly is also consistent with increased myo-II contractile activity (Blanchoin et al., 2014; Haviv et al., 2008; Wilson et al., 2010). Thus, both stress fiber assembly and loss are possible, even within a single cell, in response to increased myosin contractility (e.g. see Fig. 8A). Expression of different steady state localized patterns driven by myosin self-organization are expected to be highly influenced by mechanical context, based on in vitro studies which infer important roles for factors such as localized availability of actin cross-linking proteins, and network architecture (Backouche et al., 2006; Koenderink and Paluch, 2018; Reymann et al., 2012). These parameters, in addition to the magnitude of contractile forces experienced by local cytoskeletal structures, are likely to dictate whether persistent actomyosin assembly versus dissolution is realized over different length and time scales.

By what mechanism does loss of LINC module proteins cause increased myo-II contractile behaviors? We did not observe LMNA depletion to be associated with significant global increases to p-MLC levels in either HeLa or RPE-1 cells, though localized gain of p-MLC may be pertinent. Myosin-II activation has been best characterized in response to phosphorylation, however, increased myosin contractile activity in the absence of greater MLC (S19) phosphorylation is not unprecedented (Giuliano et al., 1992; Obara et al., 1995). Indeed, our results parallel those of a previous report which also suggested greater cell contractile activity, but without increased p-MLC levels, for cells depleted of the LINC module protein nesprin-1(Chancellor et al., 2010). We propose that altered myo-II contractile activity in cells deficient of LINC module proteins is mediated by perturbation of a feed-back regulatory cascade involving both mechanical and transcriptional control of myo-II, as illustrated (Fig. 8E).

Mechanical control of myosin function, likely through the effects of load in modifying myosin-actin catch bond behavior, is well supported by computational models and studies from cellular systems (Greenberg et al., 2016; Kasza and Zallen, 2011; Levayer and Lecuit, 2012). For instance, applied forces have been shown to control myo-II assembly and activity during fly development (Fernandez-Gonzalez et al., 2009). Our previous work and those of others suggest that the cytoplasmic actomyosin network generates tensile forces on the nucleus (Arsenovic et al., 2016; Wiggan et al., 2017). We suggest that obligate resisting forces provided by the nucleus, via LINC module mechano-coupling, provide important mechanical feedback signaling cues to help control the contractile activity of myo-II across the cytoplasmic actomyosin network. LINC module disruption is proposed to impair physical mechano-coupling and feedback that likely entails the activity of the nuclear envelope protein emerin, which contributes to tuning of nuclear mechanical resistance (Guilluy et al., 2014). Further studies utilizing more rapid and acute procedures for nuclear envelope disruption will be required to examine these initial myo-II activation events. Nevertheless, a model for the nucleus playing a general role in mechano-regulation of the cytoplasmic cytoskeletal network is supported by recent data (Ambrosini et al., 2019). Distinct roles for individual Sun and nesprin LINC complex proteins, non-actomyosin cytoskeletal elements, and chromatin remodeling are feasible (Hamouda et al., 2020).

We further propose that increased contractile activity and actomyosin assembly, pursuant to mechano-activation of myo-II, instigates nuclear translocation of mechanosensitive transcription factors such as MKL1 and YAP, as we observe for LMNA deficient cells in this study. Our results for YAP are similar to those shown for human myoblasts harboring mutant LMNA (Bertrand et al., 2014). Our finding that myo2A and β-actin transcript levels increased following LMNA silencing is consistent with activation of MKL1/SRF signaling, as both genes are transcriptional targets of MKL1/SRF (Esnault et al., 2014; Foster et al., 2017). Likewise, transcripts for both β-actin and myo2A were recently shown to be upregulated in cardiomyocytes from a human laminopathic model (Bertero et al., 2019), and in yet another laminopathic model, increased β-actin transcript levels coupled to aberrant MKL1-SRF activation has been observed (Osmanagic-Myers et al., 2019). Altogether, this body of work strongly support deregulated MKL1/SRF signaling as a commonality across models of LMNA deficiency. Our results (Fig. 5A) and those of others (Willer and Carroll, 2017), nonetheless, suggest complex and differential regulation of MKL1/SRF gene targets in LMNA deficient cells, as some genes such as VCL are repressed at steady state and not activated. Repression of some SRF targets like VCL in the face of increased MKL signaling may involve other gene specific regulatory factors. In preliminary studies we have found that LMNA depleted cells show increased nuclear targeting of the transcription factor YY1, which also shuttles between the nucleus and cytoplasm dependent on actin organization, analogous to MKL1 and YAP (Ellis et al., 2002). YY1 acts predominantly as a transcriptional repressor and is known to antagonize expression of some SRF regulated genes via overlapping gene promoter binding elements (Chen and Schwartz, 1997). Interestingly, the proximal promoter of VCL contains both SRF and YY1 binding sites, suggesting this hypothesis as a plausible mechanism.

The aforementioned evidence suggesting increased MKL1 activation in LMNA deficient cells is not necessarily in conflict with an initial report by Ho et al. (2013b), which advocated abrogated MKL1 signaling. Combined results from other reports suggest impairment of mechanosensitive transcription factor signaling in LINC module disrupted cells may be contextual to factors such as cell and extracellular matrix mechanical properties (Bertrand et al., 2014; Willer and Carroll, 2017). For example, impaired MKL1 and YAP nuclear translocation may be anticipated in mechanical contexts that favor steady states of self-organized actomyosin network disassembly.

In LMNA disrupted cells, induction of mechanosensitive transcription programs, such as for MKL1 and YAP, ostensibly generates a critical positive feedback loop contributing to the increased myo2A expression and assembly we uncovered. Interestingly, enucleated cell cytoplasts show impaired responses to mechanical cues and are less contractile than intact cells (Graham et al., 2018). These data support the notion that the presence of the nucleus may be relevant not only for direct mechanical feedback to actomyosin but also for gene transcription towards sustained contractile actomyosin reinforcement.

In accord with our model that deregulated actomyosin assembly is a key feature of LMNA deficiency, we identified that loss of LMNA resulted in ectopic myo2A accumulation to dynamic cell membrane structures, in the form of podosome-type adhesions and actin comet tails. Previous studies have postulated possible roles for myo-II in both podosome assembly and disassembly (Bhuwania et al., 2012), our live cell image analyses implicate roles for myo-II contractile activity in promoting F-actin polymerization at both types of membrane structures. Increased membrane-associated F-actin assembly may occur through activation of lipid bound actin assembly factors brought together by myosin induced clustering, as previously characterized (Koster et al., 2016; Vogel et al., 2017). Formation of podosome-like adhesions following LMNA loss in HeLa cells and not RPE-1 cells suggests deregulated myosin activity may contribute to different tissue specific phenotypes observed across the pleotropic traits observed for laminopathies. Intriguingly, in human pathologies, myosin-II plays an outsized role in one of the predominant tissues affected by LMNA deficiency, that being muscle. Further, we speculate that deviant membrane associated actomyosin remodeling provokes, for example, altered cellular features like podosome formation in normally competent cell types, such as osteoclasts. Modulation of podosome assembly, sites of extracelluar matrix degradation, may contribute to tissue specific laminopathic defects such as osteolysis.

In conclusion, our work establishes deregulation of myosin-II as a key mechanistic factor to cytoskeletal anomalies arising from disruption of nuclear LMNA. Myosin-II and its regulatory cascade appear attractive candidates for further therapeutic investigations towards treatment of nuclear envelopathies.

## Materials and Methods

### Cells

RPE-1, HeLa (Kyoto strain), U2OS and GM-2149 (normal fibroblasts) were grown in DMEM supplemented with 10% fetal bovine serum (Atlas Biologicals).

### Transfections

Cells were transfected with 40-50 nM of siRNAs at the time of plating and again 24 h later with either Lipofecatamine RNAiMax or Lipofectamine 2000 (Invitrogen) using manufacturer’s protocols.

### siRNA

siRNA oligonucleotides targeted to human Sun1, Sun2, nesprin1, nesprin2, lamin A/C and a control siRNA targeted to luciferase (GL2), as described (Wiggan et al., 2017), were obtained from Qiagen. siRNAs for individual targets produced similar phenotypes and were used interchangeably between experiments.

### Antibodies

Sun1, Sun2, Nesprin-1, Nesprin-2, myosin-2A, Flag-M2, alpha tubulin (Sigma); p-myosin light chain (Ser19 and Thr18/Ser19), (Cell Signaling); p-MLC (immunoblots, gift of R. Wysolmerski); lamin A/C (sc-7292), MKL1 (c-19), MKL1 (H-140), YAP (sc-101199), MLC (sc-4814) (Santa Cruz); GAPDH (Millipore); ADF/cofilin (1439, (Shaw et al., 2004).

### Gene expression analysis

Total mRNA was isolated using the RNeasy Mini Kit (Qiagen) and cDNA prepared using the iScript cDNA synthesis kit (Bio-rad) following the manufacturer’s procedures. Quantitative PCR (qPCR) was performed in duplicate or triplicate using the iQ SYBR green reagent with a Bio-rad CFX 96 thermocycler. Target gene expression was normalized to GAPDH and measured using the ΔCt method. Primers used for qPCR are listed in Table S1.

### Gene editing

Single-guide RNAs (sgRNAs) targeting N- and C-terminal coding exons of MYH9 were designed using CCTop (Stemmer et al., 2015). Guide RNAs were cloned into the plasmid pX330-U6-Chimeric_BB-CBh-hSpCas9 (Addgene plasmid 42230, (Cong et al., 2013) using procedures as described (Bauer et al., 2015). Homology directed repair donor plasmids were generated using a combination of gene synthesized DNA sequences for ∼ 225 bp homology arms and either mAID-mCherry2 derived from plasmids pMK292 (Addgene plasmid 72830) for C-terminal tagging, or sfGFP for N-terminal tagging, as described (Natsume et al., 2016). Cells, were transfected with plasmids for sgRNA and donor repair using Lipofectamine 2000. Following neomycin selection, mCherry positive cells were confirmed for proper recombination by genomic DNA PCR genotyping, as described (Bauer et al., 2015) and by visual evaluation of expected mCherry-protein cellular distribution through fluorescence microscopy. Confirmed mCherry expressing cells were subjected to a second round of CRISPR editing with plasmids for N-terminal sfGFP tagging. Dual GFP/mCherry positive cells were isolated by fluorescence activated cell sorting and characterized as described earlier. Sequences for sgRNAs, homology arms and PCR primers for genotyping are listed in Tables S2-3.

### Cell culture, immunofluorescence staining and microscopy

Fluorescent images were acquired with an Olympus IX81 spinning disk confocal (CSU22 head) microscope with either 100x/1.40 NA, 60x/1.42 NA or 40x/1.35 NA objectives. Images were acquired with a Photometrics Cascade II CCD camera using SlideBook (Intelligent Imaging Innovations). Some live cell images were captured in TIRF mode on a Nikon Eclipse TiE inverted microscope with 100x/1.40 NA to an Andor iXon Ultra 888 EMCCD or Clara camera.

For live cell imaging, cells plated on glass bottom 35 mm dishes were housed in a stage incubator at 37°C with CO_2_ in regular growth medium. Cells were co-transfected with siRNAs and/or plasmids encoding either GFP-Lifeact (Riedl et al., 2008), pCMV-Flag-MKL1-N100 (derived from Addgene plasmid 19848), or FLAG-MKL1 or S449A/T450A/S454A (Addgene plasmid 19845, Muehlich et al., 2008). Phenotypic analyses of siRNA treated cells were at 48-72 h post initial siRNA treatment. In serum stimulation experiments cells were starved for 24h in medium containing 0.3% serum and stimulated by replacing with medium containing 15% serum. Cells were treated with 100 μM blebbistatin (Sigma) or with DMSO for controls, in experiments assessing myosin-II inhibition. For fixed cell microscopy, cells grown on glass coverslips were fixed in 4% formaldehyde in CBS buffer (10 mM MES pH 6.1, 138 mM KCl, 3 mM MgCl_2_, 2 mM EGTA, 0.32 M sucrose) for 20 min at room temperature. For cytoskeletal staining 0.4% Triton X-100 was included in the fix buffer. Fluorescently labeled phalloidin (F-actin labeling), Alexa-594 conjugated DNaseI (G-actin labeling) and secondary antibodies were from Invitrogen. DNA was labeled by 4’,6-diamidino-2-phenylindole (DAPI). Confocal z-stacks were acquired for analyses of both live and fixed cells.

### Image and data analysis

Quantification of fluorescence intensity and image data analyses were performed using CellProfiler (Carpenter et al., 2006), ImageJ, Python and open source Python packages including Pyqtgraph (www.pyqtgraph.org) and Pandas. Focal adhesions were segmented and quantified by first applying a Difference of Gaussian filter to basal confocal fluorescence images of paxillin labeling. Adhesions, from all cells in each captured field, were then segmented by the triangle thresholding method. Intensity measurements were made from background subtracted images on adhesions of size ≥ 0.5 μm^2^. For nuclear/cytoplasmic measurements, projection images from three nuclear midpoint planes based on DAPI labeled confocal z-stacks captured at 0.2 - 0.3 μm steps were utilized. Intensity values for each experiment were generally normalized to range between .01 and 1. Maximum intensity projection images of the entire z-stack were utilized for whole cell measurements. G/F-actin ratios were measured from cells dual labeled with phalloidin and DNase I. Images were processed by a CellProfiler pipeline, as described (Wiggan et al., 2017), for segmentation and data extraction. Myosin-II filament intensities were measured from line profile peak intensities of fluorescence images of C-terminal mCherry labeled Myo2A puncta, along stress fibers. Microtubule and myosin filament anisotropy was measured using the ImageJ plugin, FibrilTool, from mid-cell level confocal z-stack planes of randomly selected ∼ 6 μm^2^ cell regions of myo2A or tubulin immuonstained images, as detailed (Boudaoud et al., 2014). Live cell fluorescence images of cells expressing Myo2A-mCh and GFP-Lifeact were enhanced by a Laplacian of Gaussian filter. Super-resolution radial fluctuations (SRRF) images were generated using the default settings of the ImageJ plugin, NanoJ-SRRF (Gustafsson et al., 2016), from 100 confocal frames captured at 10 −250 ms exposures.

F-actin abnormality, based on comparison to normal control cells, was assessed as follows: cells with evident increased numbers of basal stress fibers (SF) that were maintained over the central region of the cells were classified as ‘increased SF’; cells with much reduced levels or complete absence of basal stress fibers were classified as ‘decreased SF’; cells with a mixture of SF organizations often characterized by a disrupted network of stress fibers, the presence of actomyosin asters, F-actin condensates, fragmented F-actin bundles and F-actin bundles with frayed ends were classified as ‘disrupted SF’. Mean Pearson’s correlation coefficients and 95% confidence intervals were calculated following Fisher’s Z-transformation. Statistical analyses were conducted using either Python or R statistical packages. Data were tabulated from three or more experiments with the exception of Figures 2E, and 5C (Starved and 1h serum stimulation) which were from two experiments. Where appropriate either a two-sided Welch’s t-test, or one-way ANOVA followed by either Tukey-Kramer or Dunnett’s post hoc (for comparisons to control) tests were utilized.

### Supplemental Material

Figure S1 shows heterogeneous cytoskeletal defects within and between cells following LMNA depletion. Figure S2 shows actomyosin aster organization in RPE-1 cells. Figure S3 shows ectopic expression of active MKL1 fails to attenuate F-actin cytoskeletal disorders resulting from LMNA silencing. Figure S4 shows live cell analysis of myosin-II inhibition and recovery. Table S1 lists primers for qPCR. Table S2 lists CRISPR editing related sequences. Table S3 lists CRISPR genotyping PCR primers. Video S1 shows actomyosin dynamics at podosome-like adhesions. Video S2 shows F-actin comet formations in LMNA depleted RPE-1 cells. Video S3 shows actomyosin organization during comet formation in HeLa cells. Video S4 shows myosin-2A stress fiber dynamics in response to blebbistatin treatments.

## Acknowledgements

We thank Jennifer Skinner for assistance with experiments, and Dr. R. Wysolmerski for sharing antibodies. Supported in part by a CSU Core Infrastructure Grant for Microscopy, a Boettcher Foundation Webb-Waring Biomedical Research Program award (TSJ), a grant from the Pew Biomedical Scholars Program (JGD), NIH grants NS064217 (OW), GM088371 (JGD), R35GM119728 (TSJ), R01AG049668 and AG044812 (JRB).

## Author Contributions

Conceptualization, O.W, T.S.J, J.R.B.; Investigation O.W.; Resources, O.W., J.G.D, T.S.J, J.R.B.; Writing – Original Draft, O.W., J.G.D, T.S.J, J.R.B.; Writing – Review & Editing, O.W., J.G.D, T.S.J, J.R.B.; Funding Acquisition, O.W., J.G.D, T.S.J and J.R.B.

## Supplementary Material Legends

**Figure S1.**
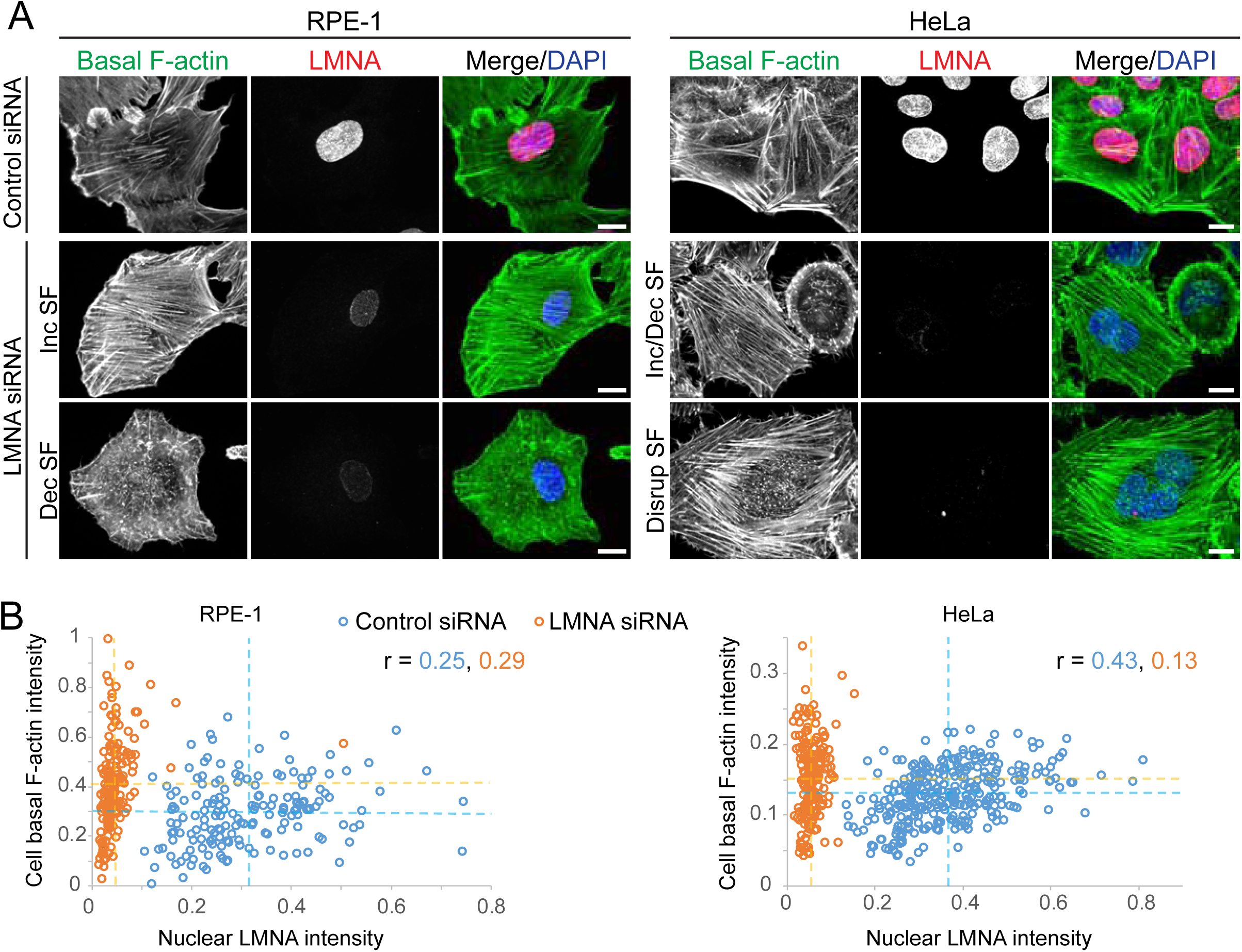
Heterogeneous cytoskeletal defects within and between cells following LMNA depletion. (A and B) Confocal immunofluorescence images and quantification of basal F-actin features, illustrating no appreciable correlation between residual nuclear LMNA levels and divergent F-actin phenotypes. Horizontal and vertical lines (B) depict mean values, r is Pearson’s correlation coefficient. Scale bars, 10 μm.

**Figure S2.**
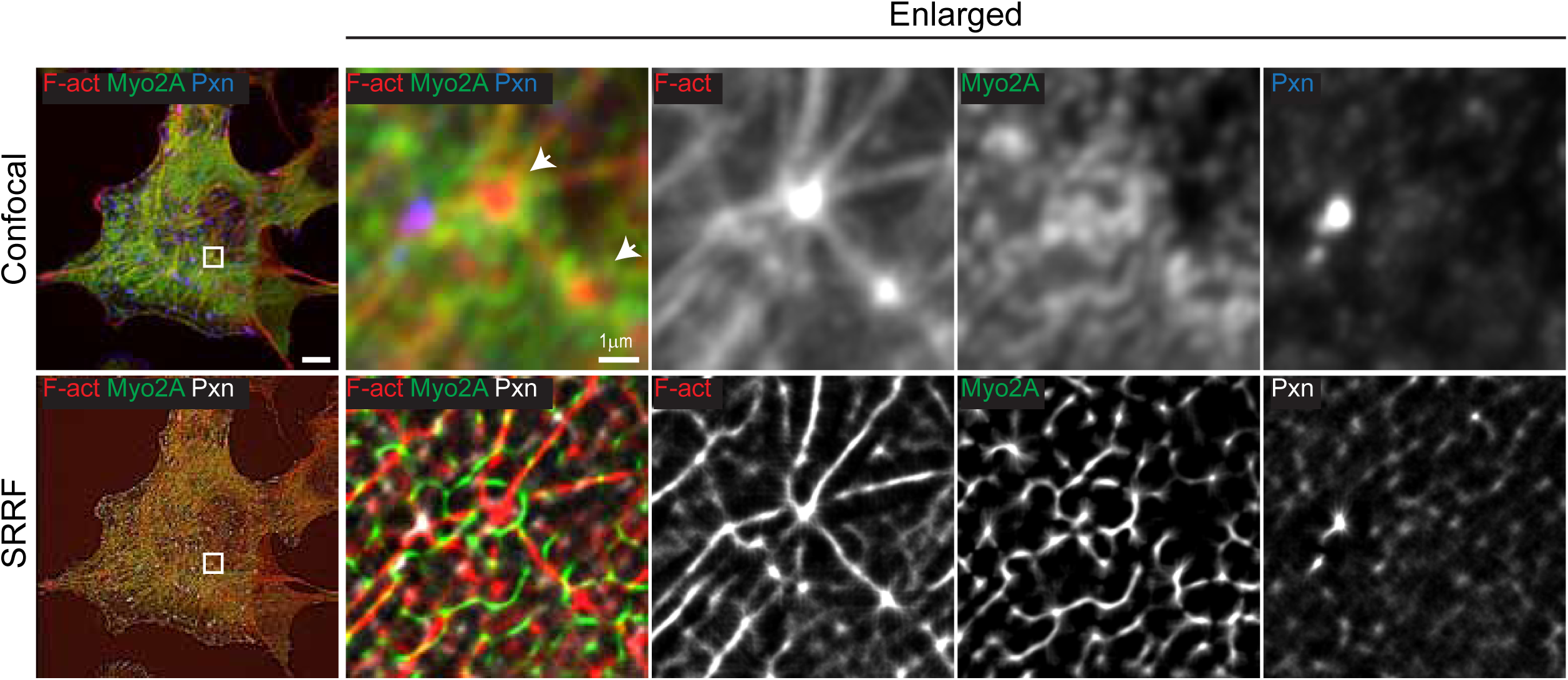
Actomyosin asters in RPE-1 cells lack podosome vinculin organization. Confocal immunofluorescence (upper panels) and SRRF super-resolved images of LMNA siRNA treated RPE-1 cells labeled for paxillin (pxn), F-actin (F-act) and myosin-2A (myo2A). Boxed regions (left column) are shown as enlargements. Arrows depict actin foci surrounded by myosin rings but absence of vinculin rings. Scale bar, 10 μm or as indicated.

**Figure S3.**
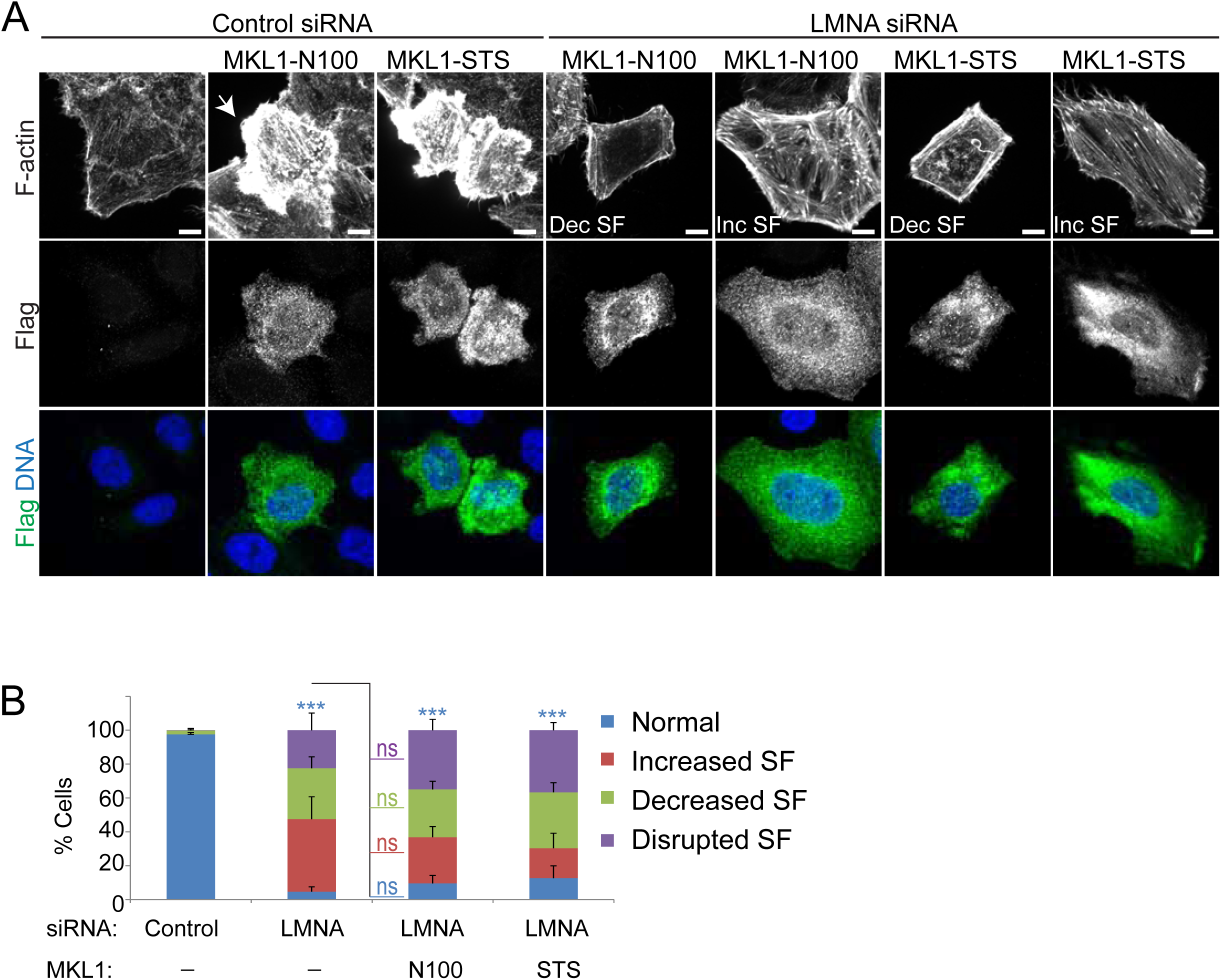
Ectopic expression of active MKL1 fails to attenuate F-actin cytoskeletal disorders resulting from LMNA silencing. (A) Confocal immunofluorescence images of HeLa cells at 72h post siRNA treatments and 48h post transfection of vectors expressing Flag tagged MKL1 mutant proteins. (B) Quantification of F-actin phenotypes. Values are mean ± SD, n ≥ 341/treatment. *** p < .001, one-way Anova and Dunnett’s post hoc test for normal phenotype relative to control; ns, not significant, Tukey’s post hoc test.

**Figure S4.**
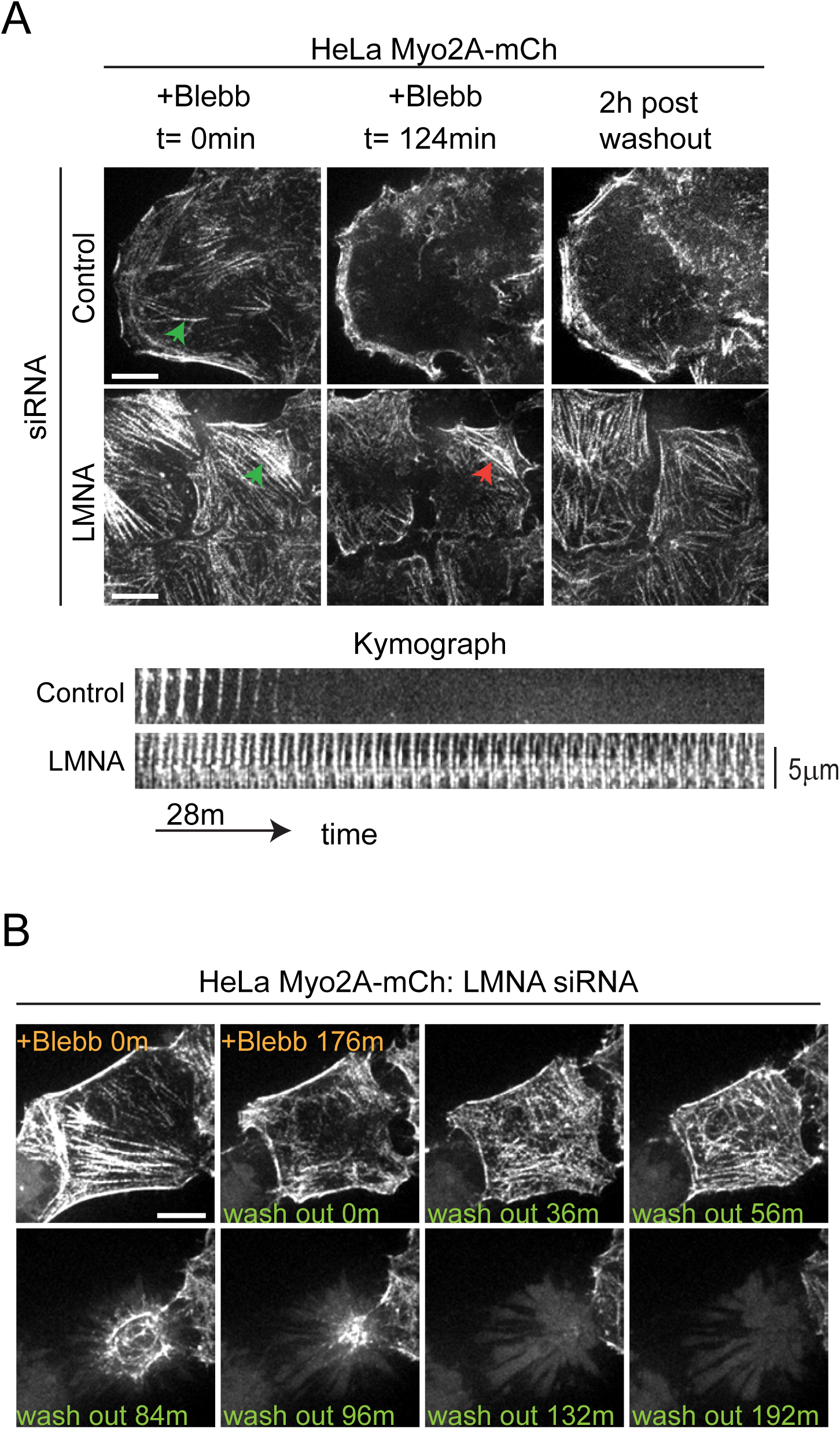
Live cell analysis of myosin-II inhibition and recovery. (A and B) Fluorescence time-lapse images of blebbistatin treatment (+Blebb) and drug washout. Green arrowheads depict stress fibers from which kymographs (A) were generated. Red arrow highlights stress fiber persistence following blebb treatment. Scale bars, 10 μm.

**Video S1**. Time-lapse confocal microscopy of basal z-projection planes for HeLa cells expressing myosin-2A-mCherry and transiently transfected GFP-Lifeact (F-actin labeling). Images were captured at 1 min intervals, 48h post siRNA treatment and the video rate is 20 frames/s. Related to Figure 3.

**Video S2**. Time-lapse confocal microscopy of control and LMNA siRNA depleted RPE-1 cells depicting F-actin (GFP-Lifeact) comet formations (red circles). Images were captured at 5 sec intervals and the video rate is 10 frames/s. Related to Figure 4.

**Video S3**. Time-lapse confocal microscopy of basal z-projections for LMNA siRNA treated HeLa cells expressing myosin-2A-mCherry and transiently transfected GFP-Lifeact. Red arrow illustrates comet tail with myosin-2A labeling. Images were captured at 90 sec intervals and the video rate is 3 frames/s. Related to Figure 4.

**Video S4**. Time-lapse confocal microscopy of basal z-projections for control or LMNA siRNA treated HeLa cells expressing myosin-2A-mCherry. Cells were treated with 100 μM blebbistatin and washed as indicated. Images were captured at 7 min intervals and the video rate is 20 frames/s. Related to Figure 8.

